# Preferred auditory temporal processing regimes and auditory-motor interactions

**DOI:** 10.1101/2020.11.14.382051

**Authors:** Pius Kern, M. Florencia Assaneo, Dominik Endres, David Poeppel, Johanna M. Rimmele

## Abstract

Decoding the rich temporal dynamics of complex sounds such as speech is constrained by the underlying neuronal processing mechanisms. Oscillatory theories suggest the existence of one optimal perceptual performance regime at auditory stimulation rates in the delta to theta range (<10 Hz), but reduced performance in the alpha range (10-14 Hz) is controversial. Additionally, the widely discussed motor system contribution to timing remains unclear. We measured rate discrimination thresholds between 4-15 Hz, and auditory-motor coupling strength was estimated through auditory-motor synchronization. In a Bayesian model comparison, high auditory-motor synchronizers showed a larger range of constant optimal temporal judgments than low synchronizers, with performance decreasing in the alpha range. This evidence for optimal auditory processing in the theta range is consistent with preferred oscillatory regimes in auditory cortex that compartmentalize stimulus encoding and processing. The findings suggest, remarkably, that increased auditory-motor interaction might extend such an optimal range towards faster rates.

---

Natural sounds such as speech or music contain temporal structure at multiple time scales. Particularly slow acoustic modulations in the delta-theta range (2-9 Hz) are considered crucial for speech and music processing (Ding et al., 2017; Pellegrino et al., 2011; Singh & Theunissen, 2003). Such natural statistics are arguably not accidental and co-occur with potential neuronal coding principles in auditory cortex (Ravignani et al., 2019; Singh & Theunissen, 2003). Support for this proposal comes from electrophysiological studies that identified endogenous oscillations in auditory cortex in the delta-theta band (Giraud et al., 2007; Keitel & Gross, 2016; Lakatos et al., 2005; Lubinus et al., 2019). By entraining to acoustic signals at these time scales, neuronal oscillations in auditory cortex might contribute to the processing of temporal information in sound (Ghitza, 2012; Giraud & Poeppel, 2012; Gross et al., 2013; McAuley & Jones, 2003; Miller & McAuley, 2005; Rimmele, Gross, et al., 2018). The optimal processing range of neuronal populations should, therefore, constrain auditory perception, by facilitating auditory temporal processing within this range (Haegens & Zion Golumbic, 2018; Rimmele, Morillon, et al., 2018).

Perceptual constraints, such as decreased neuronal tracking of speech and reduced speech comprehension at fast rates outside of the presumably optimal range, have been shown previously (Ahissar et al., 2001; Brungart et al., 2007; Doelling et al., 2014; Ghitza & Greenberg, 2009). Similarly, for amplitude modulated sounds and isochronous tone sequences, reduced neuronal tracking (Teng et al., 2017; Teng & Poeppel, 2020) and reduced temporal sensitivity have been observed at stimulus rates associated with the higher alpha range compared to lower rates (Drake & Botte, 1993; Friberg & Sundberg, 1995; Teng et al., 2017; Viemeister, 1979). Overall, there is considerable evidence from studies on rate perception for ‘constant’ optimal auditory temporal processing in the theta range (4–8 Hz, including lower rates in the delta range 2-4 Hz). This has typically been assessed through relative difference thresholds for rate discrimination, i.e. the minimal difference between two stimulation rates necessary for discrimination normalized by the standard rate. Relative difference thresholds have been shown to be lowest and constant in the theta range, which is commonly interpreted as a zone of optimal temporal processing, referring to Weber’s law (Drake & Botte, 1993; Friberg & Sundberg, 1995; Viemeister, 1979). According to Weber’s law, the ability to distinguish stimulation rates is proportional to the frequency of the presentation rate. The absolute rate difference necessary for discrimination, thus, scales with the stimulation rate, resulting in a constant relative difference threshold. Although, the temporal sensitivity has been shown to be constant at low stimulation rates (corresponding roughly to the delta-theta band, in neural terms), the onset of the decrease in temporal sensitivity is controversial. While some studies already find higher relative thresholds for rates around 8 Hz (Drake & Botte, 1993; Friberg & Sundberg, 1995; ten Hoopen et al., 1994; Ten Hoopen et al., 2011), others report a threshold increase at 10 Hz (McAuley & Kidd, 1998; Michon, 1964), 12 Hz (Ehrlé & Samson, 2005; Elliott & Theunissen, 2009), 16 Hz (Nordmark, 1968; Viemeister, 1979), or even 40 Hz (Dau et al., 1997; Sheft & Yost, 1990). In these studies, typically only a coarse range of standard (modulation) rates in the upper theta and alpha range was tested.

Here we investigate whether interindividual differences in auditory-motor coupling strength might contribute to the controversial or mixed findings regarding the (onset of) sensitivity changes. ‘Temporal predictions’ from motor cortex have been shown to modulate auditory processing in studies presenting periodic tone sequences (Arnal et al., 2015; Morillon et al., 2014; Morillon & Baillet, 2017) or continuous speech (Keitel et al., 2018; Park et al., 2015), even during passive listening (Chen et al., 2008; Grahn & Rowe, 2013). Recently, Assaneo et al. (2019) developed a simple behavioral protocol to measure spontaneous auditory-motor synchronization. The synchronization of one’s own speech production to a perceived speech stream differed widely but systematically across individuals, revealing a bimodal distribution of high and low synchronizers. On a neuronal level, high synchronizers showed stronger functional and structural connectivity between frontal speech-motor and auditory cortices, rendering this behavioral protocol suitable to estimate individual differences in auditory-motor coupling strength. Furthermore, speech production more strongly modulated perception in high synchronizers compared to lows, indicating increased motor top-down predictions with increasing auditory-motor coupling strength (Assaneo, Ripollés, et al., 2019; Assaneo et al., 2020). Furthermore, recently it has been proposed based on MEG findings that the right hemispheric lateralization of speech processing is reduced at more challenging fast stimulation rates (Assaneo, Rimmele, et al., 2019), whereas such lateralization might be linked to motor top-down predictions (Tang et al., 2020).

We investigate the following hypotheses: (1) There is a particular auditory sensitivity for processing stimulation rates in the theta range, that decreases at higher rates in the alpha range, reflected in increasing rate discrimination thresholds. (2) Interindividual differences in auditory-motor coupling strength, estimated with the spontaneous speech synchronization test (SSS-test) (Assaneo, Ripollés, et al., 2019), modulate auditory temporal processing sensitivity. Based on previous findings, we expect (a) an overall increased auditory temporal sensitivity in high compared to low synchronizers due to increased temporal top-down predictions from the motor cortex. (b) Alternatively, the benefit might particularly occur at higher “non optimal” rates. In an adaptive weighted up-down staircase procedure, we tested auditory discrimination thresholds for a fine-grained range of rates in the theta to alpha range (4 - 15 Hz). Bayesian model comparison was used to test our hypotheses. To validate the threshold measure, we used the method of constant stimuli to measure rate discrimination performance at two rates in the theta and alpha range (4, 11.86 Hz).

## Method

### Participants

All participants reported normal hearing and absence of any type of dyslexia, neurological or psychiatric disorder or intake of psychotropic substances during the last six months. The experimental procedures were ethically approved by the Ethics Council of the Max Planck Society (No. 2017_12). All participants gave written informed consent prior to the study and received monetary compensation. Participants were excluded from the statistical analyses because of outlier behavioral performance (*n* = 3; mean relative difference threshold in at least one psychophysical procedure exceeded the median +/−3 median absolute deviation; Leys et al., 2013) or because the two runs in the SSS-test were inconsistent (*n* = 2). The final sample included 55 participants (age range: 19–32 years (*M* = 25.38, *SD* = 3.78), 27 females, 3 left-handed), clustered into 35 high and 20 low synchronizers.

### Procedure

The experiment was conducted in a sound attenuated experiment booth. Stimulus presentation and response acquisition were run on a Windows 7 computer using the Psychophysics Toolbox (Brainard, 1997; Pelli, 1997) for Matlab R2017a (version 9.2). Responses were collected on a standard computer keyboard. Auditory stimuli were presented binaurally at a 44100 Hz sampling rate via a RME Fireface UCX audio interface, a Phone Amp Lake People G109 headphone amplifier and Beyerdynamic DT770 Pro (80 Ohm) closed-back headphones. In the SSS-test, we used Etymotic Research 3C insert earphones (50 Ohm) with foam ear tips and a directional gooseneck condenser microphone (Shure MX418 Microflex) placed about 5 cm in front of the participant for speech recording.

### Rate discrimination task

We adopted an auditory two-interval forced-choice (2IFC) rate discrimination task (Fig. 1a), in which a standard sequence and a faster comparison were presented in random order (Drake & Botte, 1993). Participants were asked to indicate as accurately and quickly as possible by keypress whether the first or second sequence was faster. They were instructed to guess whenever they failed to discriminate the rate and to refrain from any movement to the beat such as tapping.

**Fig. 1.**
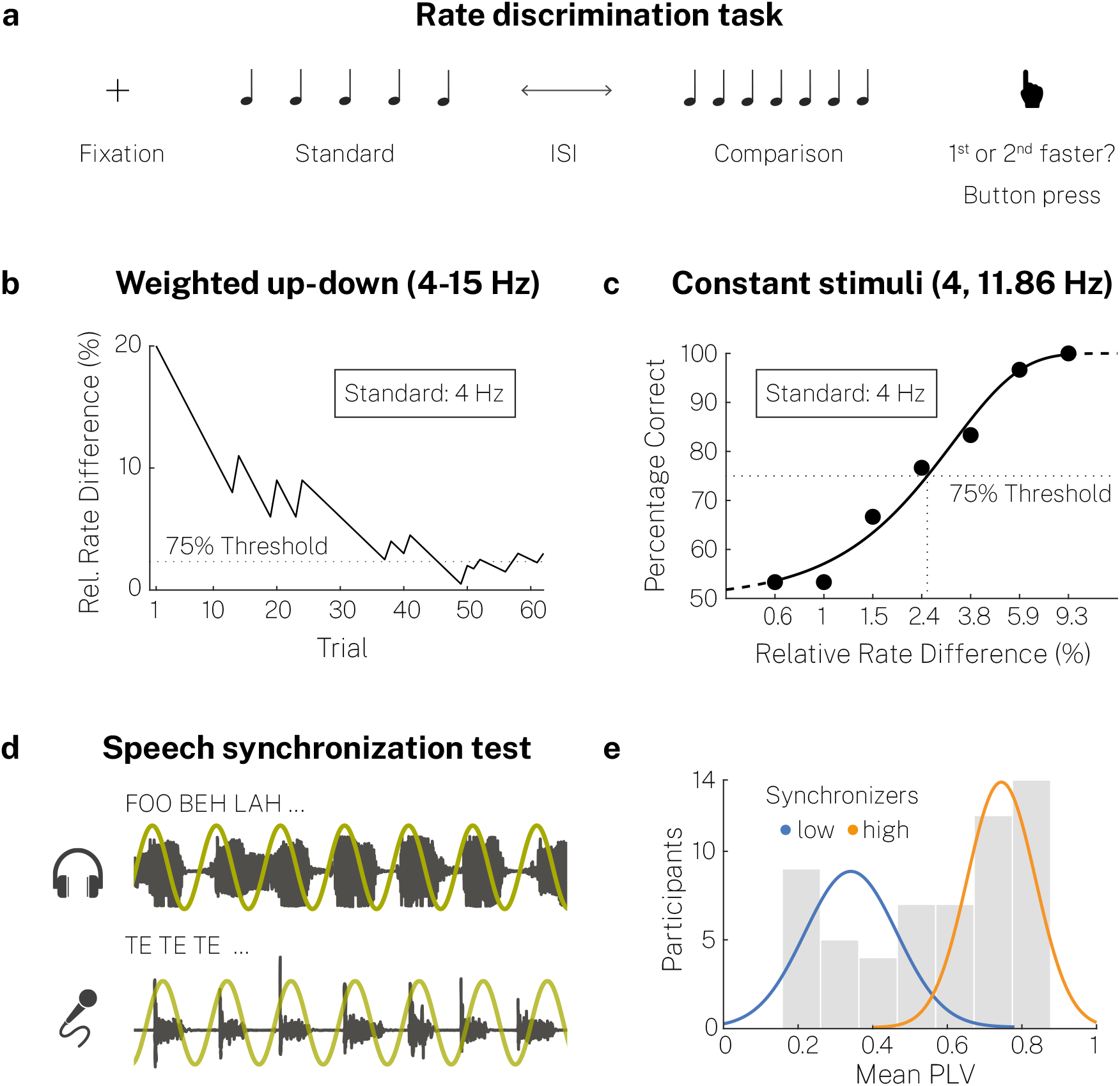
Experimental design. (a) In a 2-IFC task, participants judged whether the first or second of two isochronous tone sequences was faster. Standard and comparison sequence were randomly presented at either position. (b) Using a weighted up-down staircase procedure, we measured relative difference thresholds at eight standard rates between 4 and 15 Hz (Exemplary run from one participant at a standard rate of 11.86 Hz). (b) Additionally, the discrimination performance was measured in a constant stimuli procedure for two standard rates at 7 stimulus levels (i.e. levels of the comparison sequence). The psychometric function was fitted to estimate the relative difference threshold (Exemplary data and psychometric function from one participant at a standard rate of 4 Hz). (d) In the auditory-motor speech synchronization test, participants whispered the syllable /te/ while listening to a random syllable train in two runs (Assaneo, Ripollés, et al., 2019). The synchronization between the produced and presented speech was measured (icons based on resources from Flaticon.com). (e) According to the mean phase-locking value (PLV), we clustered participants into two groups, low and high synchronizers.

We generated isochronous sequences by concatenating a 15 ms 440 Hz sinusoidal pure tone (5 ms rise/fall of the sound envelope) with silent tone intervals, depending on the stimulation rate. One of the two sequences consisted of 5 and the other of 7 tones, randomized trial-by-trial. Auditory stimuli were presented at 70 dB SPL (peak amplitude normalization).

On each trial, a fixation cross appeared 500–750 ms prior to stimulus onset (random uniform distribution). Standard and comparison sequences were presented in random order separated by an inter-sequence interval (random uniform distribution between 4.5 and 5.5 times the tone inter-onset interval of the first sequence). After giving their response, participants received visual feedback (correct, incorrect) in the weighted up-down procedure, but not in the method of constant stimuli. The intertrial interval was 1000–1500 ms (random uniform distribution).

### Weighted up-down procedure

We used an adaptive weighted up-down (WUD) staircase method implemented in the Palamedes toolbox (Kingdom & Prins, 2010) to measure discrimination thresholds block-wise for eight standard rates linearly spaced from 4 to 15 Hz (Fig. 1b). The relative difference between standard and comparison rate was reduced by a certain step size given a correct response and was increased by three times that step size following an incorrect response to converge on a 75% correct level, defined as the threshold (Kaernbach, 1991). All parameters were chosen in terms of the relative difference between standard and comparison rate (in Hz), given by *d*_*r*_ = (*rate*_*comparison*_ – *rate*_*standard*_) / *rate*_*standard*_. Start values for each standard rate condition (linearly increasing across rates from 20–27%) were selected well above the expected threshold to observe a convergence to threshold level and to capture the threshold area in all participants at each standard rate (Green, 1990) based on previous studies and pilot results. We chose linearly increasing start values to avoid differential adaptation and learning effects as higher thresholds were expected at higher rates. The initial step size (1%) was divided in half after six (0.5%) and again after twelve reversals (0.25%), i.e. after twelve changes of direction in the adaptive run (e.g., up to down). Progressively decreasing step sizes were expected to yield reliable and fine-grained threshold estimates more efficiently (Levitt, 1971; Rammsayer, 1992). In case the relative rate difference became smaller or equal to one step-size, the current difference was divided in half after a correct response.

The relative difference threshold for each standard rate was computed as the mean of the last 6 reversals. Short breaks were included every 20 trials. One adaptive run ended after 18 reversals. To familiarize participants with the task, a minimum of 5 and a maximum of 15 training trials (75% correct criterion) were completed at the beginning of each block. In training trials, the relative difference between standard and comparison rate was 10% higher compared to the start values in the main part (i.e. 30–37%, 1% steps).

### Method of constant stimuli procedure

We used the method of constant stimuli (CS) to validate the WUD threshold estimates by fitting the psychometric function at two standard rates of main interest, 4 and 11.86 Hz (Fig. 1c). The psychometric function describes the relationship between a participant’s response behavior and a stimulus feature (Wichmann & Hill, 2001). We measured performance (percentage correct) at seven comparison levels (relative rate difference in the comparison sequence). For every participant, we calculated the rate of the comparison sequences by multiplying a seven-point scale logarithmically spaced from 0.2 to 3 with the individual WUD threshold to capture the whole range of the psychometric function from 50% chance level to 100% correct (see: Herbst & Obleser, 2019). Every run contained one trial of each comparison level in random order (mixed presentation). There were 30 runs in total, amounting to 30 trials measured per comparison level. Pauses were included after 21 trials. The procedure was completed for each standard rate separately in randomized order across participants.

We fitted a Weibull function to the individual data at both standard rates using Bayesian inference methods implemented in the Psignifit Toolbox 4 (Schütt et al., 2016). The guess rate was fixed at 50% and three parameters were estimated: the 75% correct threshold, the width, and the lapse rate. We assessed the goodness-of-fit for individual data by comparing the deviance to a parametric bootstrap sample distribution (*N* = 10000, 95% percentile).

### Auditory-motor speech synchronization test

In the SSS-test (Fig. 1d; for details see Assaneo, Ripollés, et al., 2019) we measured participants’ ability to synchronize their speech production to a heard syllable sequence. First, participants wearing headphones were instructed to cautiously increase the volume of a background babble while continuously whispering the syllable /te/, until they could not hear their own speech anymore (start sound level: 70 dB SPL, maximum: 95 dB SPL). In two training trials, participants listened to a 10 s periodic syllable /te/ sequence (at 4.5 Hz) and were asked to whisper /te/ at the same rate for 10 s directly afterwards. Finally, a 70 s random syllable train with a progressively increasing rate (*M* = 4.5 Hz, range: 4.3–4.7 Hz, steps: 0.1 Hz after 60 syllables) was presented. The random syllable stream audio file was created using the MBROLA synthesizer (male American English voice, 200 Hz pitch) (Dutoit et al., 1996). Participants were instructed to whisper /te/ continuously in synchrony with the audio. Two blocks were measured for each participant.

We quantified auditory-motor speech synchronization by calculating the phase locking value (PLV) (Lachaux et al., 1999) between the cochlear envelope of the audio stimulus and the envelope of the recorded speech signal (time windows 5 s, overlap 2 s). The cochlear envelope was determined using the NSL Auditory Model toolbox for Matlab (auditory channels: 180–7246 Hz). Phases of both envelopes were calculated via the Hilbert transform after resampling to 100 Hz and band-pass filtering (3.5–5.5 Hz). To classify high and low synchronizers, we performed k-means clustering (Arthur & Vassilvitskii, 2007) of participants into two clusters based on mean PLVs across both blocks (Fig. 1e, mean PLV across high synchronizers = 0.74, *SD* = 0.1, mean PLV across low synchronizers = 0.34, *SD* = 0.12). Note that one participant was not assigned to one cluster consistently given the nature of the clustering algorithm. Our findings were not altered by the classification of this participant as high synchronizer (reported here, because this was the case in ~58% of 10000 simulation runs), as low synchronizer, or exclusion.

Finally, participants filled out a questionnaire on demographic data, task strategies, and musical sophistication (Goldsmiths Musical Sophistication Index, Gold-MSI, Müllensiefen et al., 2014).

### Analysis

Data analysis was performed in Matlab 2018b (version 9.5) and in R (version 3.6.1). We performed a Bayesian model comparison using the RStan software (version 2.19.2) (Carpenter et al., 2017) to specifically test our hypotheses on temporal sensitivity in the WUD procedure depending on stimulation rate and auditory-motor speech synchronization behavior. We compared 34 models *M*_*i*_, *i* ∈ {1, 2, …, 34}, each specified by a set of parameters *θ* (Fig. 2a). For model simplicity, we assumed that the observations *D* (relative difference thresholds) are independent and identically distributed given the models’ parameters. We approximated the data generation process by log-normal distributions, given that threshold values could only be positive and were expectedly right-skewed (Gaussian normal distributions truncated at zero yielded lower posterior probabilities, while the main results were equivalent). We further assumed that the variance in each standard rate condition and subgroup (high and low synchronizers) was drawn from the same distribution.

**Fig. 2.**
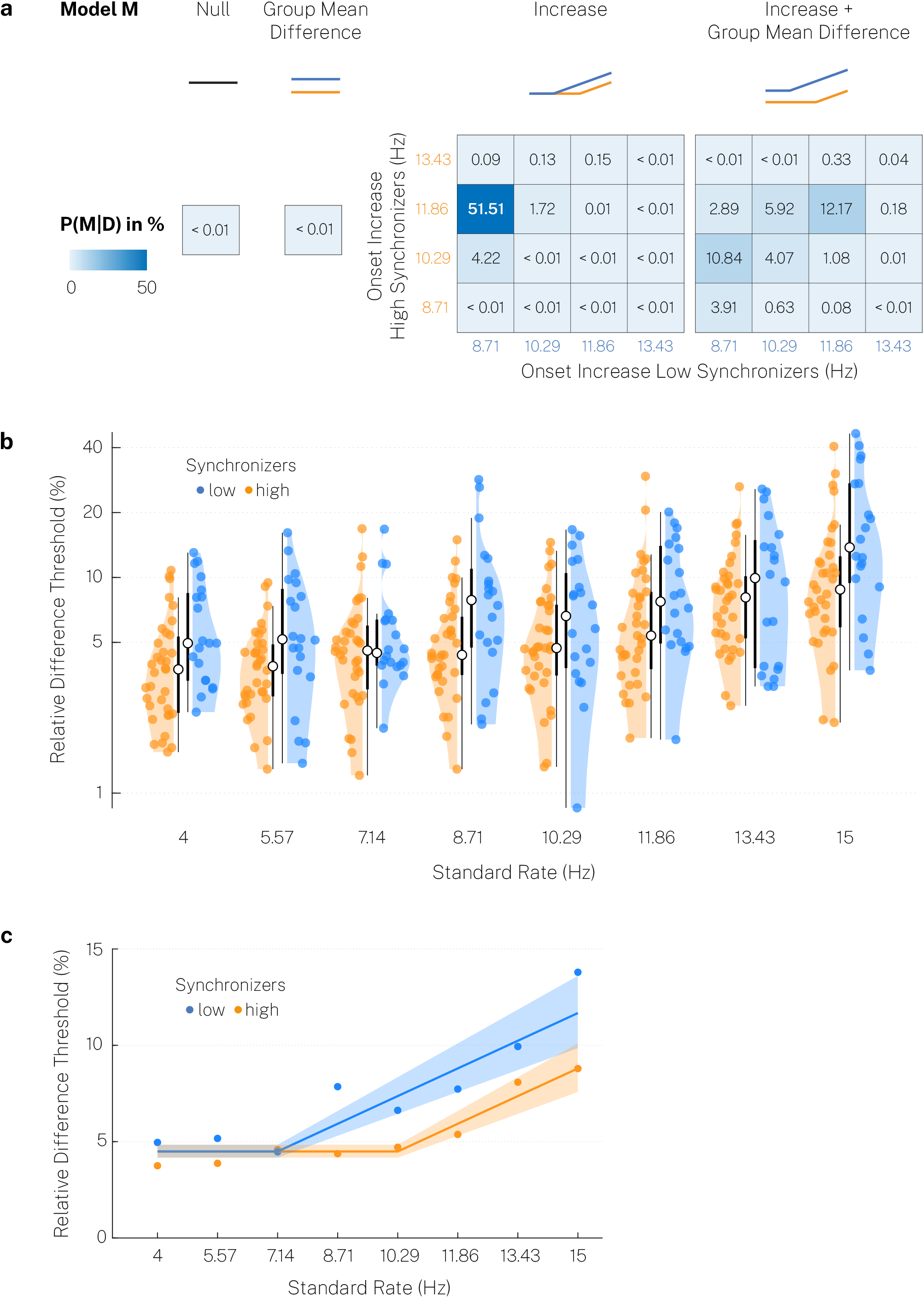
Weighted up-down method indicates an optimal processing range modulated by auditory-motor coupling. (a) We tested 34 models (first row) that included either a constant threshold (null model), a difference between high and low synchronizers (group mean difference), a linear threshold increase at different starting points that could vary between groups (increase), or both (increase + group mean difference) and calculated the posterior probability of each model given the data, P(M|D) (last row). (b) Relative difference thresholds for high and low synchronizers. Colored dot: individual participant, white dot: median, thick line: quartiles, thin line: quartiles ± 1.5 × interquartile range. (c) Median relative difference thresholds (dots) and model predictions of the best model (line) with 95% confidence interval (shaded area) for high and low synchronizers.

In the null model *M*_*1*_, the relative difference threshold was constant at *θ*_1_ across all standard rates.

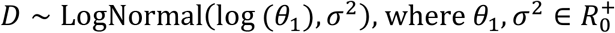

The prior distribution for *θ*_1_ was set to be a normal distribution with prior mean *θ*_1−0_ and prior variance 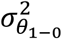. Summarizing previous results on rate discrimination in isochronous sequences (Drake & Botte, 1993; Ehrlé & Samson, 2005; Friberg & Sundberg, 1995; ten Hoopen et al., 1994; Ten Hoopen et al., 2011), we expected the mean threshold *θ*_1−0_ across the tested frequency range to be 5% on average, with single observations ranging from minimally 0.1% to maximally 50%. We doubled the maximal variance possible given that range to obtain a prior estimate for 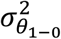. The same prior distribution for *θ*_1_ was used in the remaining models as well.

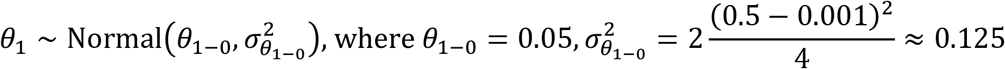

The prior distribution for *σ*^2^ was specified as a uniform distribution with a lower bound *α* = 0. The upper bound *β* was determined by the maximum expected variance, given an expected minimal threshold of 0.1% and a maximum threshold of 50%. The same prior distribution for *σ*^2^ was used in the remaining models as well.

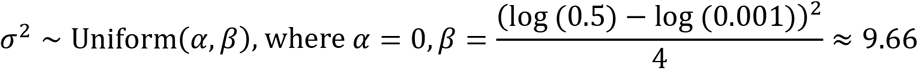

Model *M*_*2*_ included the group difference in relative difference thresholds *θ*_2_ between high and low synchronizers, coded by the indicator variable *x*_2_. We constrained *θ*_2_ to be greater than zero as we predicted that low synchronizers would have higher threshold values compared to high synchronizers.

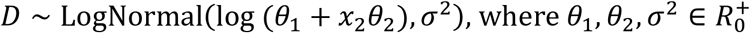

Based on differences between musicians and non-musicians in previous studies, we expected the threshold to be on average 4% higher in low synchronizers and set the prior mean *θ*_2−0_ of the Gaussian prior correspondingly. The prior variance 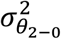 was set equal to 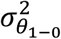 since we assumed that the variance in group differences would not be higher than in the overall mean. The same prior distribution of *θ*_2−0_ was applied in the remaining models.

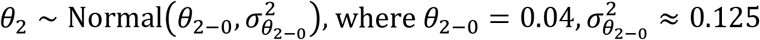

In models 3 to 18, we included the hypothesized departure from a constant relative threshold *θ*_1_ in the theta range and model a linear threshold increase in the alpha range by including a slope parameter *θ*_3_. We considered four different starting points of the linear increase (8.71, 10.29, 11.86, and 13.43 Hz), coded by indicator variables (e.g. *x*_1_ = 0,0,0,1,2,3,4,5}). The starting point could vary in high versus low synchronizers, which was permuted in the 16 different models.

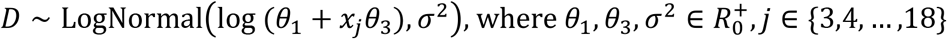

The prior distribution of *θ*_3_ was a normal distribution with a prior mean *θ*_3−0_ of 2% predicated upon previous results on threshold increases in the alpha range. The prior variance 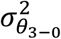 was set equal to again 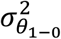. The same prior distribution was adopted in the remaining models as well.

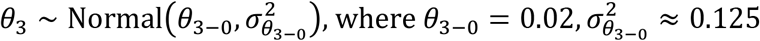

The models 19 to 34 included both effects, a constant difference between high and low synchronizers and a linear threshold increase in the alpha range with varying starting points.

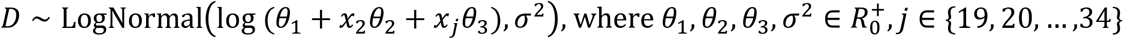

We assigned all models equal prior probability, 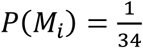. According to Bayes’ rule, the posterior probability of a model M_i_ is given by the product of its prior probability and the marginal likelihood of the observations given the model divided by the marginal probability of the observations:

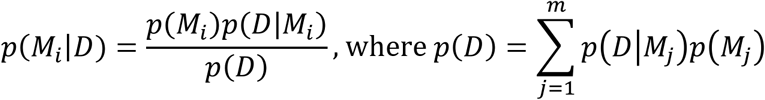

The Bayes Factor was calculated to determine the amount of evidence in favor of one model as the ratio between its posterior probability and the posterior probability of the remaining models (Kass & Raftery, 1995). Similarly, we obtained the probability for a specific effect by marginalizing over all secondary effects and comparing the resulting posterior probability.

We used numerical methods implemented in the RStan and the bridgesampling package (Gronau et al., 2017) in R to estimate the log posterior probability. We first obtained samples of the posterior distribution (log density function) by running each model in RStan, which uses a No-U-Turn (NUTS) sampling Markov chain Monte Carlo (MCMC) algorithm (Carpenter et al., 2017). We ran 5 chains (maximal tree depth = 10, target average acceptance probability for adaptation = 0.95), each containing 15000 iterations, including 5000 warm-up iterations, to receive 50000 sample draw iterations in total. Further increasing the number of iterations did not change the parameter estimates and posterior probabilities, indicating convergence. The accuracy of the MCMC algorithm was checked via the diagnostics provided in the RStan environment (rhat statistics, divergences, saturation of the maximum three depth, and the Bayesian fraction of missing information). Finally, we used the RStan output to compute the log marginal likelihood via bridge sampling and, based on that, the posterior probability (Gronau et al., 2017).

## Results

### Weighted up-down procedure

Median relative difference thresholds in the weighted up-down (WUD) procedure ranged from 4.08% at 4 Hz to 10.5% at 15 Hz (Threshold averaged across standard rates in each participant: *Mdn* = 5.25%, *MAD* = 2.54%). High synchronizers displayed lower thresholds averaged across all standard rates (*Mdn* = 5.52%, *MAD* = 2.15%) compared to low synchronizers (*Mdn* = 8.21%, *MAD* = 3.7%) (Fig. 2b; Wilcoxon rank-sum test, one-sided, *W* = −2.25, *p* = .012, *r* = −0.21). We used a Bayesian model comparison to test the influence of stimulation rate and auditory-motor speech synchronization behavior more specifically. The Bayesian sampling algorithm displayed appropriate behavior for all of the 34 models (rhat statistics < 1.001; no divergence in any of the fits; no saturations of the maximal tree depth 10; non-suspicious Bayesian fraction of missing information). Bridge sampling therefore allowed for accurate estimation of the posterior probability of each model given the data (Fig. 2a). As we used a uniform prior probability for a relatively large number of models 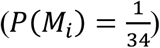, we did not expect a high Bayes factor (BF) for a single model. More importantly, the posterior probability of the most likely model (M5) increased considerably from a prior probability of about 3% to 51.51% (*BF* = 1.06). The model M5 included a constant threshold from 4 to 7.14 Hz in low synchronizers and from 4 Hz to 10.29 Hz in high synchronizers (Fig. 2c). Crucially, we additionally found moderate evidence for an earlier threshold increase in low synchronizers compared to high synchronizers, when comparing the corresponding models to all the remaining ones (*BF* = 3.51). In contrast, a general group difference between high and low synchronizes across all standard rates was not supported by our data when comparing the corresponding models to all the remaining ones (*BF* = 0.73).

### Constant stimuli procedure

To validate the WUD threshold measure procedure, we examined rate discrimination sensitivity in the constant stimuli (CS) procedure by fitting the psychometric function at 4 Hz and 11.86 Hz (Fig. 1c). Inspection of goodness-of-fit for individual data demonstrated proper fit for all 110 individual fits (55 subjects, 2 standard rates; see Fig. 3a for mean data in each standard rate condition across all participants). The lapse rate was generally low, with a trend for higher lapse rates at 11.86 Hz (*Mdn* = 1.16 × e–07%, *MAD* = 2.07%) compared to the 4 Hz standard rate condition (*Mdn* = 6.05 × e–08%, *MAD* = 1.3%) (Wilcoxon signed-rank test, two-sided, *T* = −1.93, *p* = .054). This suggests that participants remained attentive throughout the whole procedure, producing only a few lapses.

**Fig. 3.**
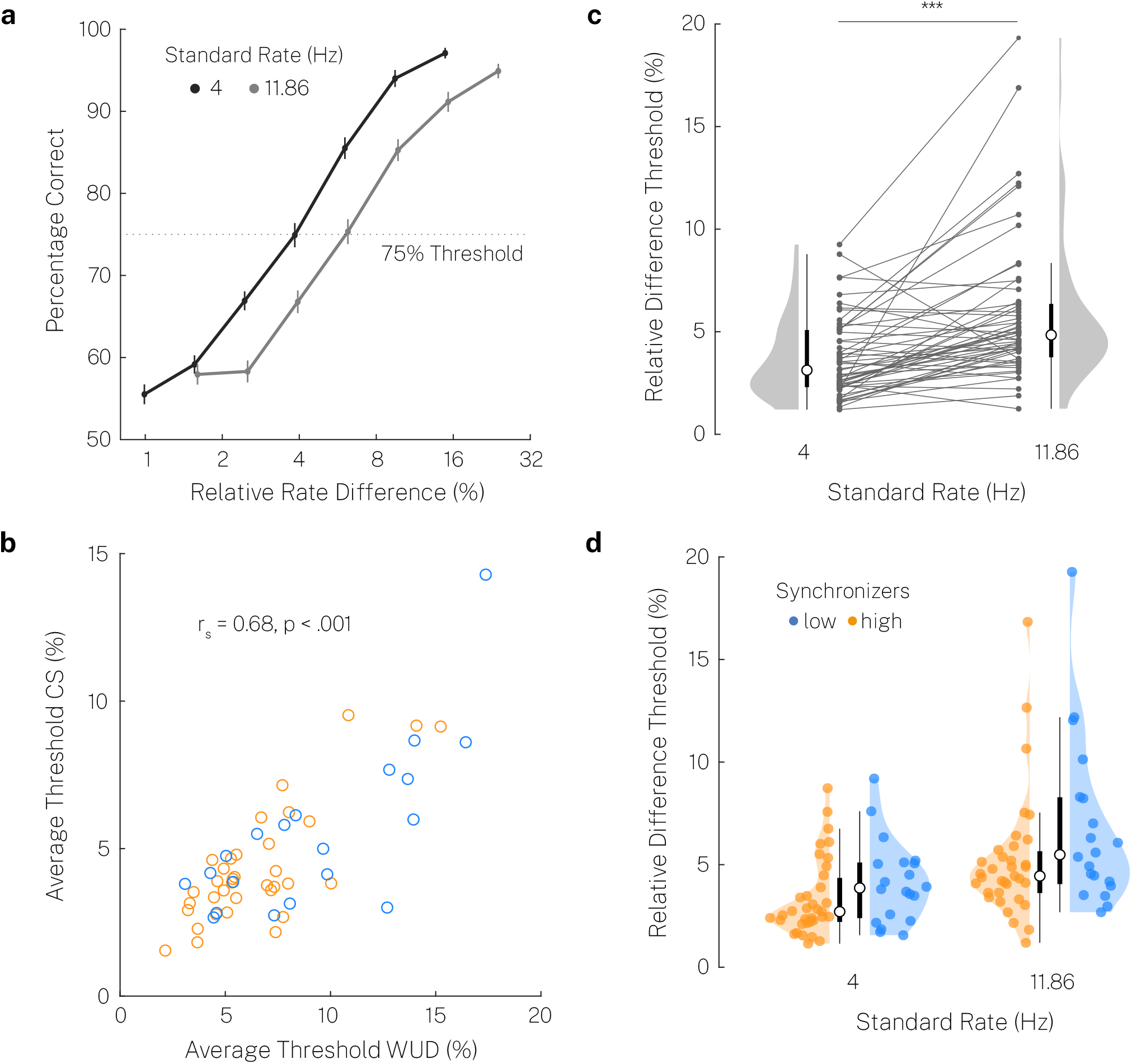
Constant stimuli method suggests optimal processing range in the theta range. (a) Mean psychometric functions across all participants at two standard rates and seven comparison levels. Error bars represent the standard error of the mean. (b) Scatter plots of average thresholds across standard rates in the WUD and CS procedure. Correlation calculated on high/low group-demeaned variables. (c) Relative difference thresholds at two standard rates across all participants. Grey dot and horizontal line: individual participant, white dot: median, thick line: quartiles, thin line: quartiles ± 1.5 × interquartile range. Thresholds were significantly lower at rates of 4 Hz compared to 11.86 Hz. (*** p < .001.) (d) Thresholds for high and low synchronizers. Colored dot: individual participant.

Relative thresholds at 4 Hz and 11.86 Hz strongly correlated between the WUD and CS method (Average across standard rates: Spearman-rank correlation, high/low synchronizer group-demeaned variables, *r*_*s*_ = 0.68, *p* < .001, Fig 3b; 4 Hz: *r*_*s*_ = 0.66, *p* < .001; 11.86 Hz: *r*_*s*_ = 0.55, *p* < .001), as well as the difference in thresholds between 4 Hz and 11.86 Hz (*r*_*s*_ = 0.39, *p* = .003). Thresholds were overall lower in the CS procedure compared to the WUD procedure at both 4 Hz (Wilcoxon signed-rank test, two-sided, *T* = −4.37, *p* < .001, *r* = −0.42) and 11.86 Hz (Wilcoxon signed-rank test, two-sided, *T* = −3.24, *p* = .001, *r* = −0.31), presumably reflecting increased familiarity with the task. But the difference between relative difference thresholds at 4 and 11.86 Hz (indicating a threshold increase at rates in the alpha range) did not differ between methods (Wilcoxon signed-rank test, two-sided, *T* = 0.91, *p* = .361). Together, these results confirm that temporal sensitivity was measured adequately in the adaptive weighted up-down procedure, which was our main procedure to test temporal sensitivity across the theta and alpha range.

Thresholds in the CS method were also higher at 11.86 Hz (*Mdn* = 4.84%, *MAD* = 2.38%) compared to 4 Hz (*Mdn* = 3.13%, *MAD* = 1.55%) (Fig. 3c; Wilcoxon signed-rank test, one-sided, *T* = 4.95, *p* < .001, *r* = 0.47). This provides further evidence for a decrease in temporal sensitivity from the theta to the alpha range at 11.29 Hz. Average thresholds across standard rate conditions were descriptively lower in high (*Mdn* = 3.82%, *MAD* = 1.42%) versus low synchronizers (*Mdn* = 4.87%, *MAD* = 2.05%), but there was only a trend for statistical significance (Wilcoxon rank-sum test, one-sided, *W* = −1.58, *p* = .057) (Fig. 3d). (Note that in the CS method we only tested two rates, and thus could not access group differences as nuanced as in the WUD threshold measure).

### Correlation of temporal sensitivity, auditory-motor coupling, and musicality

Our previous analyses suggest a close relationship between auditory-motor speech synchronization behavior and auditory temporal sensitivity. Here, we analyzed the impact of general musical sophistication (Gold-MSI) on these measures. High synchronizers reported higher musicality compared to low synchronizers (Wilcoxon rank-sum test, one-sided, *W* = 3.27, *p* < .001, *r* = 0.3). Given the bimodal distribution of auditory-motor speech synchronization behavior, we demeaned all variables in the high and low synchronizers clusters to control for spurious correlations when calculating correlations among the variables (Fig. 4). Musical sophistication correlated positively with mean PLVs in the SSS-test (*r*_*s*_ = 0.46, *p* < .001), indicating that higher musicality went along with improved auditory-motor synchronization behavior. There was only a trend for a negative correlation between musical sophistication and average relative threshold levels across standard rates in the CS procedure (*r*_*s*_ = −0.26, *p* = .056) and the WUD procedure (*r*_*s*_ = −0.23, *p* = .092). Crucially, threshold levels in the WUD method correlated moderately with mean PLVs in the SSS-test (*r*_*s*_ = −0.36, *p* = .008), even when controlled for musical sophistication (partial correlation *r*_*s*_ = −0.29, *p* = .034). This suggests that differences in auditory-motor synchronization behavior explained individual variance in temporal sensitivity beyond musical sophistication. However, the correlation between average threshold levels in the CS procedure and mean PLVs did not reach statistical significance (*r*_*s*_ = −0.21, *p* = .13).

**Fig. 4.**
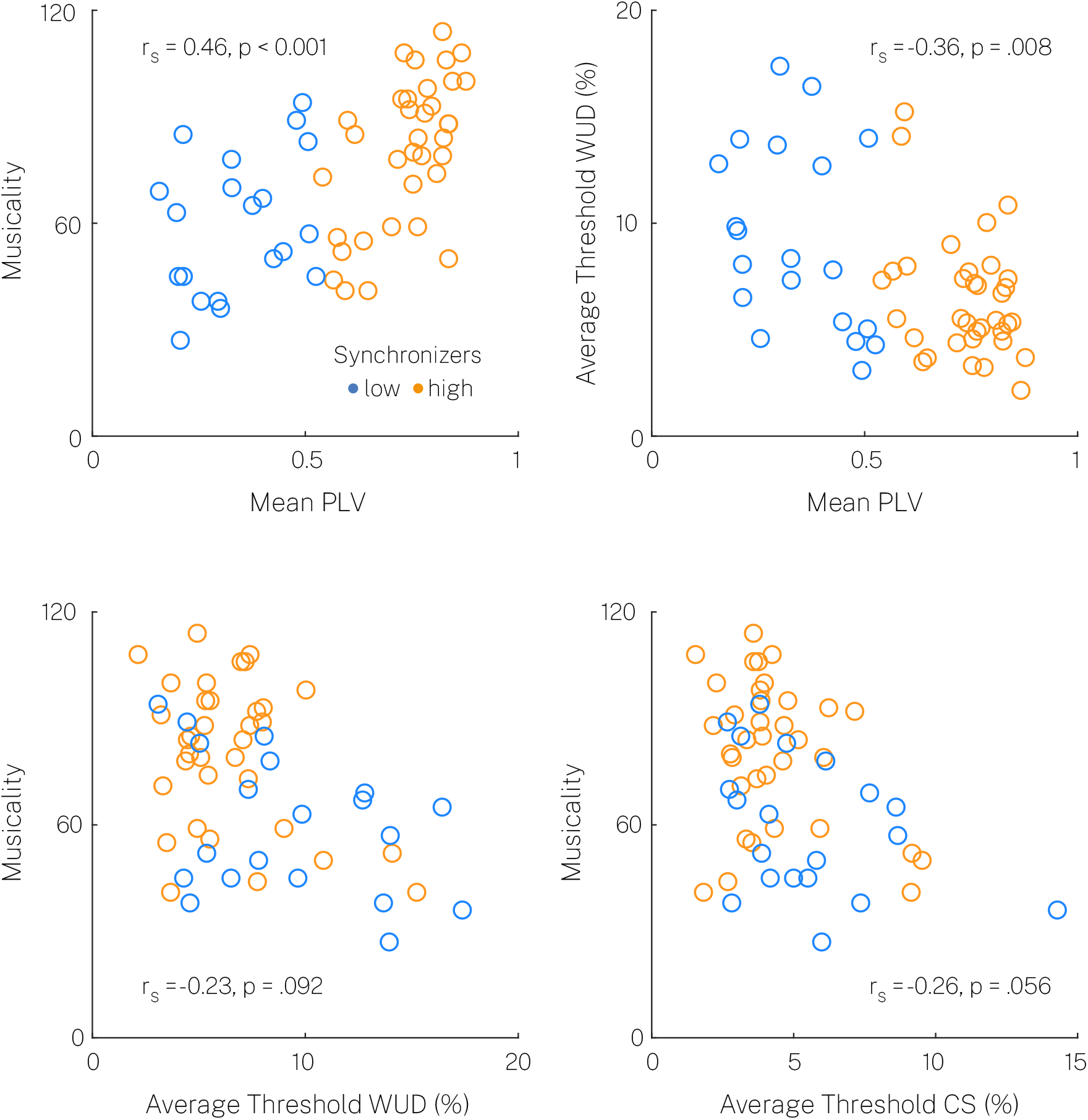
Scatter plots and correlations between rate discrimination threshold, auditory-motor coupling, and musicality measures. Correlations were calculated based on high/low group-demeaned variables to account for spurious correlations.

## Discussion

The experiments we present lie at the intersection of two fundamental questions in perception that have been investigated by and large independently. On the one hand, rate sensitivity is a basic question in auditory perception. On the other hand, the interaction between perception and action systems has been increasingly studied, with new insights about auditory-motor coupling. We combine these two lines of research in psychophysical experiments and discover a surprising generalization: listeners with increased coupling between their auditory and speech motor systems have broader sensitivity to acoustic rates. Our evidence invites two conclusions: first, there are indeed preferred, fine-grained regimes of temporal processing in hearing at low frequency modulation rates in the theta range. Second, the motor system influences perceptual timing; in particular, individual variability in auditory-motor coupling undergirds listeners temporal processing thresholds, with a larger range of optimal processing in high compared to low auditory-motor synchronizers.

Overall, individuals showed constant low relative rate discrimination thresholds in the theta range. The range of optimal processing varied for high and low synchronizers, with the most likely optimal range for low synchronizers from 4 to 7.14 Hz and high synchronizers from 4 to 10.29 Hz. Thresholds increased at frequencies above the optimal range up to 15 Hz, indicating reduced temporal sensitivity in the alpha range (and above). Some previous studies report reduced temporal sensitivity for rate discrimination at higher rates in the alpha range (Drake & Botte, 1993; Friberg & Sundberg, 1995; McAuley & Kidd, 1998; ten Hoopen et al., 1994; Ten Hoopen et al., 2011), whereas others did not (Dau et al., 1997; Elliott & Theunissen, 2009; Sheft & Yost, 1990; Viemeister, 1979). Compared to previous reports, we tested rate discrimination within a fine-grained range of rates in a larger sample of participants. Furthermore, we applied an additional procedure, the constant stimuli method, to confirm that the thresholds were reliably estimated by the adaptive weighed up-down procedure. Our data support previous behavioral findings, and neurophysiological evidence, suggesting that neuronal oscillatory activity in auditory cortex shows optimal temporal processing in the theta range (Teng et al., 2017; Teng & Poeppel, 2020). The constant relative threshold in the theta range corresponds to a linear increase in absolute rate difference required for discrimination, congruent with Weber’s law. In contrast, Weber’s law did not apply for the observed increase of relative thresholds at higher rates, suggesting a decreased temporal resolution at higher rates. In other words, this finding indicates that at higher rates the absolute rate difference required for discrimination failed to scale with the rate, i.e. the temporal resolution at higher rates seems to be constrained by a minimum ‘time constant’ which differed for high and low synchronizers (at ~7.1 ms inter-tone onset interval, or ~22.1 ms stimulus-onset interval, corresponding to ~40 Hz; see also: (Giraud, 2020; Hoonhorst et al., 2009; Joliot et al., 1994). On a neuronal level, the constant relative thresholds might reflect the preferred frequencies of the auditory cortex neuronal population.

Crucially, auditory temporal sensitivity was modulated by interindividual differences in auditory-motor coupling strength, which was estimated by testing the auditory-motor speech synchronization behavior. High compared to low synchronizers showed lower relative difference thresholds at higher rates, supporting a model that suggest a larger range of optimal temporal resolution in the high synchronizers. A possibility is that our findings reflect top-down temporal predictions from the motor system facilitating auditory processing. Previous research suggests that top-down predictions from the motor system can modulate auditory processing even during passive listening (Chen et al., 2008; Grahn & Rowe, 2013). We utilized the previously introduced behavioral spontaneous speech synchronization test (SSS-test) to distinguish individuals based on their spontaneous auditory-motor synchronization behavior (Assaneo, Ripollés, et al., 2019). The original study reported the test-retest reliability and replicability of the bimodal distribution of the synchronization behavior. Crucially, differences in functional and structural frontal-motor to auditory cortex connectivity have been shown to underly this group distinction, rendering it a suitable estimator of auditory-motor coupling strength. Furthermore, the auditory-motor coupling most likely reflects an oscillatory mechanism, whereas high synchronizers have shown stronger perceptual facilitation by top-down predictions from the motor system (Assaneo et al., 2020). Based on these previous findings, we propose that the increased rate discrimination sensitivity in high synchronizers reflects a facilitating effect of increased motor cortex recruitment in high compared to low synchronizers. Previously it has been proposed that motor top-downs effects, which are increased during demanding listening situations (e.g. such as at ‘non-optimal’ fast stimulation rates), can facilitate processing by reducing the hemispheric right lateralization resulting in bilateral auditory cortex recruitment (Assaneo, Rimmele, et al., 2019). However, whether such a neuronal mechanism can account for the observed larger optimal range for rate discrimination in high synchronizers is unknown and requires further research.

An alternative explanation of our findings is, that the differences in auditory temporal sensitivity in high and low synchronizers was due to population differences in auditory processing rather than differences in auditory-motor coupling. We cannot entirely exclude this explanation. However, previous evidence shows that the high/low synchronizer distinction comes with differences in functional and structural cortical auditory-motor interactions, rendering this “purely auditory processing” explanation unlikely. Furthermore, musical expertise might affect the temporal sensitivity for rate discrimination. For example, there is evidence for enhanced auditory-motor integration in musicians benefitting auditory processing (Du & Zatorre, 2017). Accordingly, we found that high and low synchronizers differed with respect to musical sophistication. However, the auditory-motor coupling accessed with the SSS-test (Assaneo, Ripollés, et al., 2019) correlated with inter-individual differences in temporal sensitivity beyond musicality. Our findings suggest that the SSS-test provides a more specific estimate of auditory-motor coupling compared to the (with respect to auditory-motor coupling) more indirect measure of musical sophistication.

In summary, we report a constant auditory temporal sensitivity for rate discrimination in the theta range, with a decrease in sensitivity for higher rates. Individuals with estimated strong compared to weak auditory-motor coupling showed higher rate discrimination sensitivity particularly at faster rates, indicating a larger range of optimal temporal processing. Our behavioral findings suggest constraints of auditory perception consistent with oscillatory theories, that propose neuronal populations in auditory cortex with preferred frequencies in the theta range. Furthermore, our data suggest a crucial role of auditory-motor coupling for enhancing temporal sensitivity by improving temporal resolution at higher stimulation rates.

## Acknowledgements

This work was funded by the Max-Planck-Institute for Empirical Aesthetics and supported by CLaME Max Planck NYU Center for Language Music and Emotion. The authors have no relevant financial or non-financial interests to disclose.

## Author Contributions

P. Kern and J. M. Rimmele and D. Poeppel designed the research. P. Kern collected the experimental data. P. Kern, M. F. Assaneo, D. Endres and J. M. Rimmele analyzed the data. P. Kern wrote the initial draft of the manuscript. P. Kern, M. F. Assaneo, D. Endres, D. Poeppel, and J. M. Rimmele discussed the results and contributed to the final manuscript.

## Notes

### Competing Interest Statement

The authors have declared no competing interest.

